# Genome-resolved metaproteomics decodes the microbial and viral contributions to coupled carbon and nitrogen cycling in river sediments

**DOI:** 10.1101/2022.03.11.484050

**Authors:** Josué A. Rodríguez-Ramos, Mikayla A. Borton, Bridget B. McGivern, Garrett J. Smith, Lindsey M. Solden, Michael Shaffer, Rebecca A. Daly, Samuel O. Purvine, Carrie D. Nicora, Elizabeth K. Eder, Mary Lipton, David W. Hoyt, James C. Stegen, Kelly C. Wrighton

## Abstract

Rivers have a significant role in global carbon and nitrogen cycles, serving as a nexus for nutrient transport between terrestrial and marine ecosystems. Although rivers have a small global surface area, they contribute substantially to global greenhouse gas emissions through microbially mediated processes within the river hyporheic zone. Despite this importance, microbial roles in these climatically relevant systems are mostly inferred from 16S rRNA amplicon surveys, which are not sufficiently resolved to inform biogeochemical models. To survey the metabolic potential and gene expression underpinning carbon and nitrogen biogeochemical cycling in river sediments, we collected an integrated dataset of over 30 metagenomes, metaproteomes, and paired metabolomes. We reconstructed over 500 microbial metagenome assembled genomes (MAGs), which we dereplicated into 55 unique genomes spanning 12 bacterial and archaeal phyla. We also reconstructed 2482 viral genomic contigs, which were dereplicated into 111 viral MAGs >10kb in size. As a result of integrating gene expression data with geochemical and metabolite data, we created a conceptual model that uncovers new roles for microorganisms in organic matter decomposition, carbon sequestration, nitrogen mineralization, nitrification, and denitrification. Integrated through shared resource pools of ammonium, carbon dioxide, and inorganic nitrogen we show how these metabolic pathways could ultimately contribute to carbon dioxide and nitrous oxide fluxes from hyporheic sediments. Further, by linking viral genomes to these active microbial hosts, we provide some of the first insights into viral modulation of river sediment carbon and nitrogen cycling.

**Importance:** Here we created HUM-V (Hyporheic Uncultured Microbial and Viral), an annotated microbial and viral genome catalog that captures the strain and functional diversity encoded in river sediments. Demonstrating its utility, this genomic inventory encompasses multiple representatives of the most dominant microbial and archaeal phyla reported in river sediments and provides novel viral genomes that can putatively infect these. Furthermore, we used HUM-V to recruit gene expression data to decipher the functional activities of these genomes and reconstruct their active roles in river sediment biogeochemical cycling. We show the power of genome resolved, multi-omics to uncover the organismal interactions and chemical handoffs shaping an intertwined carbon and nitrogen metabolic network and create a framework that can be extended to other river sediments. The accessible microbial and viral genomes in HUM-V will serve as a community resource to further advance more untargeted, activity-based measurements in these and related freshwater terrestrial-aquatic ecosystems.

## Introduction

The hyporheic zone (HZ) is a transitional space between river compartments, where the mixing of nutrients and organic carbon from river and groundwater stimulate microbial activity (1-3). Characterized as the permanently saturated interface between the river surface channel and underlying sediments, the HZ is considered a biogeochemical hotspot for microbial biogeochemistry (1-3), ultimately contributing to the majority of river greenhouse gas (GHG) fluxes. For instance, it is estimated rivers contribute up to 85% of inland water carbon dioxide and 30% of nitrous oxide emissions (4-6). Microorganisms in the HZ also catalyze the transformation of pollutants and natural solutes, all while microbial biomass itself supports benthic food webs(7). Together these findings highlight that microbial metabolism in HZ sediments has substantial influence on overall river biogeochemistry and health.

Despite the importance of HZ microorganisms, research linking microbial identity to specific biogeochemical reactions in the carbon and nitrogen cycles is still nascent in these sediments. In conjunction with geochemistry, microbial functional genes or gene products (e.g. *nirS* and *nrfA*) have been quantified to denote microbial contributions to specific biogeochemical pathways (e.g. nitrate reduction)(8). However, these studies often do not identify the microorganisms catalyzing the process and only focus on a few enzymatic reactions. Thus, a comprehensive assessment of the interconnected microbial metabolisms fueling carbon and nitrogen cycling in river sediments is underexplored.

More recently, 16S rRNA amplicon sequencing has shed new light on the identity of bacteria and archaeal members in river sediments. These studies revealed that cosmopolitan and dominant members in river sediments belong to six main phyla: *Acidobacteria, Actinobacteriota, Firmicutes, Nitrospirota, Proteobacteria*, and *Thaumarchaeota*(9, 10). Furthermore, in some instances cultivation paired to amplicon sequencing has assigned some of these microorganisms (e.g. *Proteobacteria*) to specific biogeochemical process (e.g., denitrification)(11). Yet, most functional inferences from taxonomic data alone are unreliable due to the dissociation between microbial taxonomy and metabolic function(12, 13). Thus, many key biogeochemical pathways in rivers (e.g., plant biomass deconstruction, denitrification, nitrogen mineralization) are not holistically interrogated alongside microbial communities(14). Furthermore, amplicon sequencing fails to sample viral communities. While it is likely viruses are key drivers of HZ microbial mortality and biogeochemical cycling by dynamics of predation and auxiliary metabolic genes, the evidence is even more sparse than for its microbial counterparts (15-18).

Cultivation-independent, community-wide, and genome resolved approaches are key to addressing the knowledge gap of how microbial and viral communities influence river biogeochemical cycling. However, metagenomic studies in river sediments are relatively sparse, and have focused primarily on gene content as opposed to whole genomes(19, 20). To our knowledge, only two river sediment studies have generated microbial genomes to allow linkages of identity to functional processes, and these have focused on the impacts of nitrate oxidizing and comammox microorganisms to nitrification(21, 22). As such, despite these recent advances the chemical exchange points that interconnect the carbon and nitrogen cycles, and the metabolic handoffs between microorganisms that sustain them cannot be discerned from existing HZ microbiome studies.

With the overarching goal of providing enhanced resolution to microbial and viral contributions to carbon and nitrogen cycling in the HZ, we created the first of its kind Hyporheic Uncultured Microbial and Viral (HUM-V) genomic catalog. We then used HUM-V to recruit metaproteomic data collected from over 30 laterally and depth distributed HZ sediment samples. We further supported this gene expression data using chemical data from paired metabolomics and geochemical measurements. Our results (i) profiled the expressed microbial metabolic exchanges that support organic and inorganic carbon and nitrogen cycling in the HZ, (ii) uncovered roles for viruses that could modulate microbial activity in the HZ, and (iii) created a roadmap of the microbial metabolic circuitry that potentially contributes to greenhouse gas fluxes from rivers. We anticipate that this publicly available community resource will advance future microbial activity-based studies in HZ sediments and is a step towards the development of biologically aware, hydro-biogeochemical predictive models.

## RESULTS and DISCUSSION

### HUM-V greatly expands the genomic sampling of HZ microbial members

We collected samples from six HZ sediment cores, with each core subsampled into six 10cm depth increments (0-60 cm). This resulted in 33 samples that were analyzed for metagenomic sequencing, geochemistry, and metaproteomics (**Fig. 1abc**). For our metagenomics data, we obtained 379Gbp of sequencing for all 33 samples which included i) the original shallow sequencing of all samples (1.7-4.9 Gbp/sample)(23) and ii) an additional deeper sequencing of 10 samples (15.3-49.2 Gbp/sample), which are reported here for the first time. We then used assembly, co-assembly, and sub-assembly approaches to reconstruct 655 bacterial and archaeal metagenome assembled genomes (MAGs). Of these genomes, 102 were denoted as medium or high-quality per the Genome Consortium Standards(24), and were dereplicated at 99% identity into the 55 unique genomic representatives that constitute the microbial component of HUM-V (see Data Availability) (**Fig. 1d**). Of the genomes retained in HUM-V, 36% were obtained from deeply sequenced, assembled, and binned samples; while 27% were from co-assemblies performed across samples. The ability to recover additional genomes relative to our prior effort which only used shallow sequencing(23) demonstrates how sequencing depth and integration of multiple assembly methods provided complementary information to generate these microbial HZ communities.

**Fig. 1.**
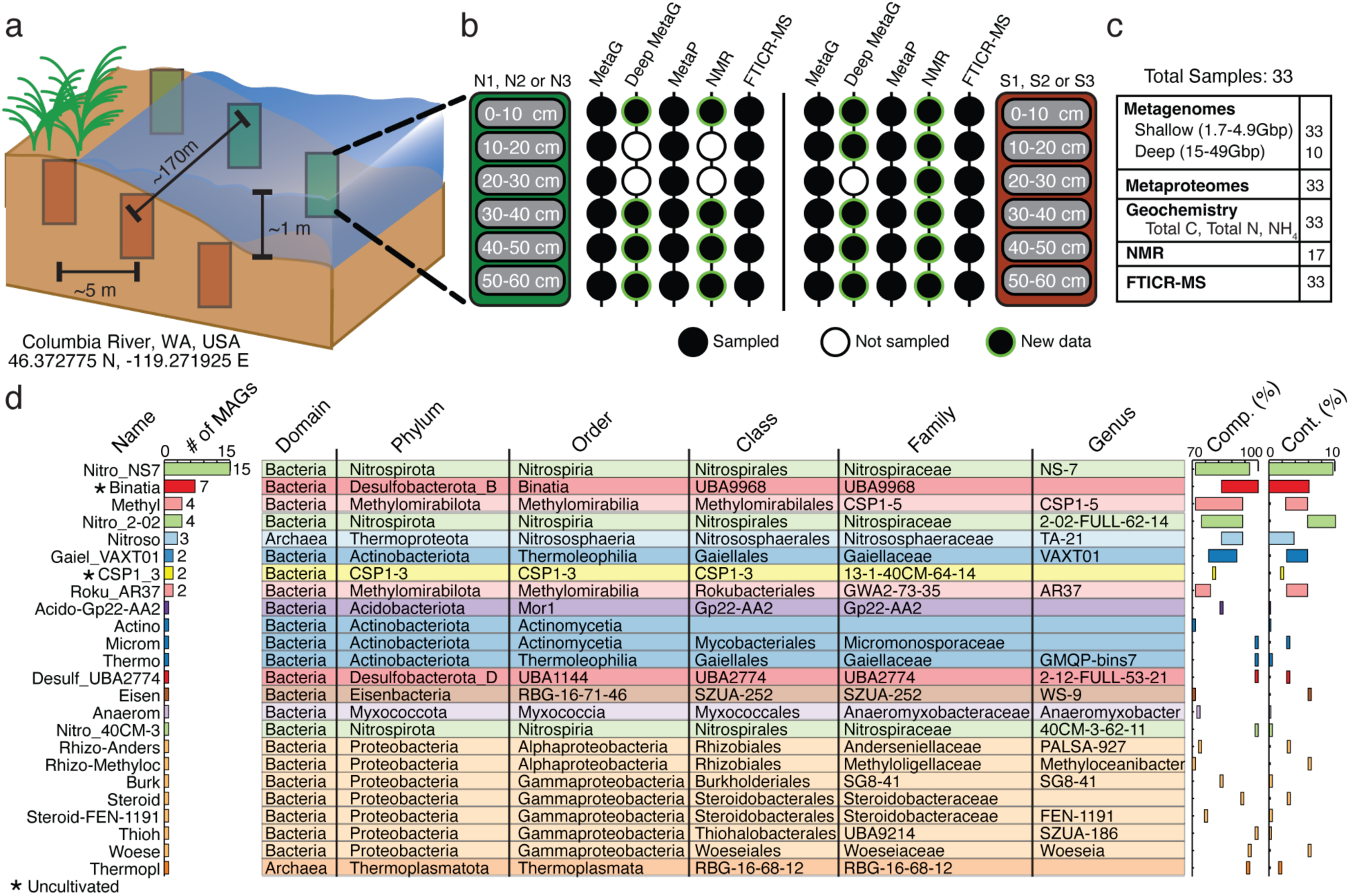
Overview of hyporheic zone sampling and the microbial genomes included in the HUM-V database. **a)** Samples were collected from two transects, each with 3 sediment cores, with each core sectioned into 10-centimeter segments from 0-60 centimeters in depth and paired with metaproteomics and geochemistry. **b)** Schematic of the data types available for each of the depth samples within a core. Black-filled circles indicate depth samples for a particular data type, open circles denote missing analyses (due to limited sample availability), and black-filled circles with green outlines indicate new sequencing that was performed as part of this project and not available in Graham et al. **c)** Summarized catalog of the total samples for each analysis. **d)** The total recovered genomes (# of MAGs), taxonomic string, inferred genome completion (Comp., %), and contamination (Cont., %) for the dereplicated microbial genomes retained in HUM-V. Asterisks on names indicate uncultivated lineages.

Given sparse metagenomic sampling of HZ sediments, it was not surprising that HUM-V contained the first genomic representatives of highly prevalent microorganisms (**Fig. 1d**, **Fig. 2ab**). Phylogenetic analyses of the 55 unique HUM-V genomes revealed they spanned 2 Archaeal and 9 Bacterial phyla, and that most genomes belonged to a subset of 3 bacterial phyla (*Desulfobacterota*, *Nitrospirota*, and *Proteobacteria*). To our knowledge, the 8 *Desulfobacterota* and 7 *Proteobacteria* genomes identified here represent the first HZ MAGs sampled from these commonly reported lineages. For the *Nitrospirota*, a prior study reported 21 MAGs that we dereplicated into 12 unique genomes (99% ANI)(21), a sampling we further expanded by an additional 20 genomes. We note the *Nitrospirota* genomes sampled here spanned 3 genera that we did not identify as being previously sampled from rivers, and that included new species within the *Nitrospiraceae 2-02-FULL-62-14*, *40CM-3-62-11*, and *NS7*. Moreover, HUM-V contains one genome of the *Actinobacteriota* that may represent a new order, as well as 6 new genera from *Acidobacteriota*, *Actinobacteriota*, *CSP1-3*, *Desulfobacterota, Proteobacteria*, and *Thermoplasmatota* (**Fig 2ab**). Further highlighting the genomic novelty of this ecosystem, HUM-V contains genomes from entirely uncultivated members of different phyla (9 genomes from *CSP1-3* and *Eisenbacteria*) and classes (10 genomes from *Binatia* and *MOR-1*). Ultimately, HUM-V is a genome resource that will enable taxonomic analyses and metabolic reconstruction of microbial metabolisms in HZ sediments.

**Fig. 2.**
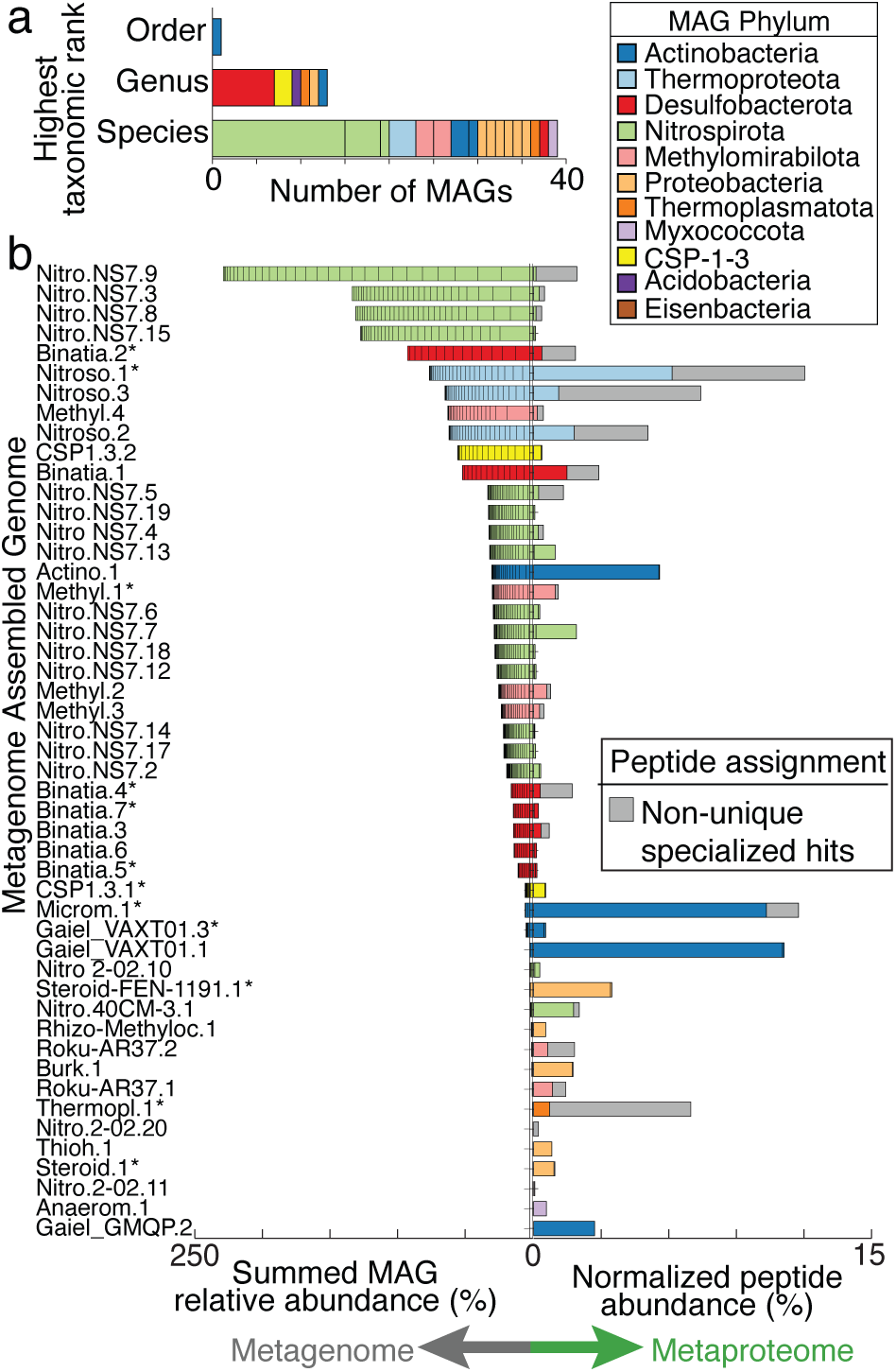
Bacterial and archaeal members of HUM-V database coupled to metaproteomics reveals active members in the hyporheic zone microbiome. **a)** Stacked bar graph indicates the taxonomic novelty of the de-replicated MAGs colored by phylum and stacked according to the first empty position within the taxonomic string provided by the Genome Taxonomy Database GTDB-Tk. Each color represents a MAG phylum according to the MAG Phylum legend, a coloring maintained across this manuscript. b) Butterfly plot reports the summed genomic relative abundance across all samples (left side) and the normalized mean proteomic relative abundance (right side) for dereplicated MAGs (55 total, 49 shown), with bars colored by phylum. MAGs that contain a partial or complete 16S rRNA sequence are denoted with and asterisk (*). Non-unique specialized peptide assignment is defined in the methods and is shown with grey bars.

### HUM-V recruits metaproteomes offering new insights into HZ microbiomes

Leveraging paired metaproteomes collected with the metagenomes allowed us to assign gene expression to each genome in HUM-V. These microbial genomes recruited 13,102 total peptides to 1,313 proteins. Because our genome analysis revealed that closely related strains shared overlapping metabolic potential (**Fig. 1d**), we analyzed the proteomic data using two approaches. First, we considered the ‘unique’ peptides that could only be assigned to proteins from a single microbial genome. These represented 67% of genes expressed in our proteome. Next, we considered proteins that recruited ‘non-unique but conserved’ peptides which we define as those assigned to proteins that (i) have identical functional annotation and (ii) are from more than one genome within the same genus. These proteins are shown in grey on **Fig. 2b**, and although they accounted for a smaller fraction of the genes expressed (14%), this prevented us excluding data due to strain overlap in our database.

In microbiome studies, dominance is often used as a proxy for microbial activity. Here, we evaluated this assumption using our paired metagenome and metaproteome data. When comparing the genome relative abundances to protein expression patterns, we observed that the most abundant genomes were not necessarily those that were most actively expressing proteins at the time of sampling. The most abundant genomes included members of the *Binatia, Nitrospiraceae NS7*, and *Nitrososphaeraceae TA-21* (formerly *Thaumarchaeota*) (**Fig. 2b**). However, only the dominant *Nitrososphaeraceae* genomes had high recruitment of the uniquely assigned proteome. On the other hand, some low abundance members (e.g., *Actinobacteriota*) accounted for a sizeable fraction (30%) of the unique proteome (**Fig. 2b**).

Leveraging these metagenomic and metaproteomic datasets, we first examined metabolic traits that were conserved across nearly all HUM-V microbial genomes. Notably, all but one of the microbial genomes recovered from this site (CSP1_3_1) encoded the genomic capacity for aerobic respiration. We defined this capability by the recovery of genes indicating a complete electron transport chain and some form of terminal oxidase (**Fig. S1**). Consistent with this genomic data, resazurin reduction assays indicated these sediments were oxygenated, and could likely support aerobic microbial respiration (**Fig. S2**). However, while proteomic evidence for aerobic respiration (cytochrome c oxidase *aa3*) was detected in nearly 40% of samples, it could only be confidently assigned non-uniquely to nitrifying *Nitrososphaeraceae.* This is likely due to the highly conserved nature of this gene, as well as the limitations of detecting membrane, heme-containing cytochromes with metaproteomic data(25). As such, we consider it likely this metabolism was more active than was captured in the metaproteomic data.

Ordination analyses of our genome-resolved metaproteomes revealed that microbial gene expression did not cluster significantly by sediment depth or transect position (**Fig. S3ab**). In fact, over 90% of measured gene expression was shared across both transects (**Fig. S3c**). When considering explanations for this lack of spatial structuring, it is possible that the microbial heterogeneity in these samples occurred over a finer spatial resolution (pore or biofilm scale, <10 cm) or larger (>60 cm) than the bulk 10 cm-depths sampled here. It is also possible that gene expression is constitutive to sustain metabolic function during highly dynamic conditions that occur in the HZ.

### An inventory of processes contributing to microbial carbon dioxide production and consumption

To uncover the microbial food web contributing to organic carbon decomposition in HZ sediments, we reconstructed the carbon degradation network using coordinated genome potential, expression, and carbon metabolite data. Based on linkages to specific substrate classes, genomes were assigned to the following different trophic levels in the carbon food chain: (i) plant polymer degradation; (ii) sugar fermentation; (iii) smaller organic compounds (e.g., alcohols and fatty acids), and (iv) single carbon compounds (carbon monoxide, carbon dioxide, methane) (**Fig. 3**).

**Fig. 3.**
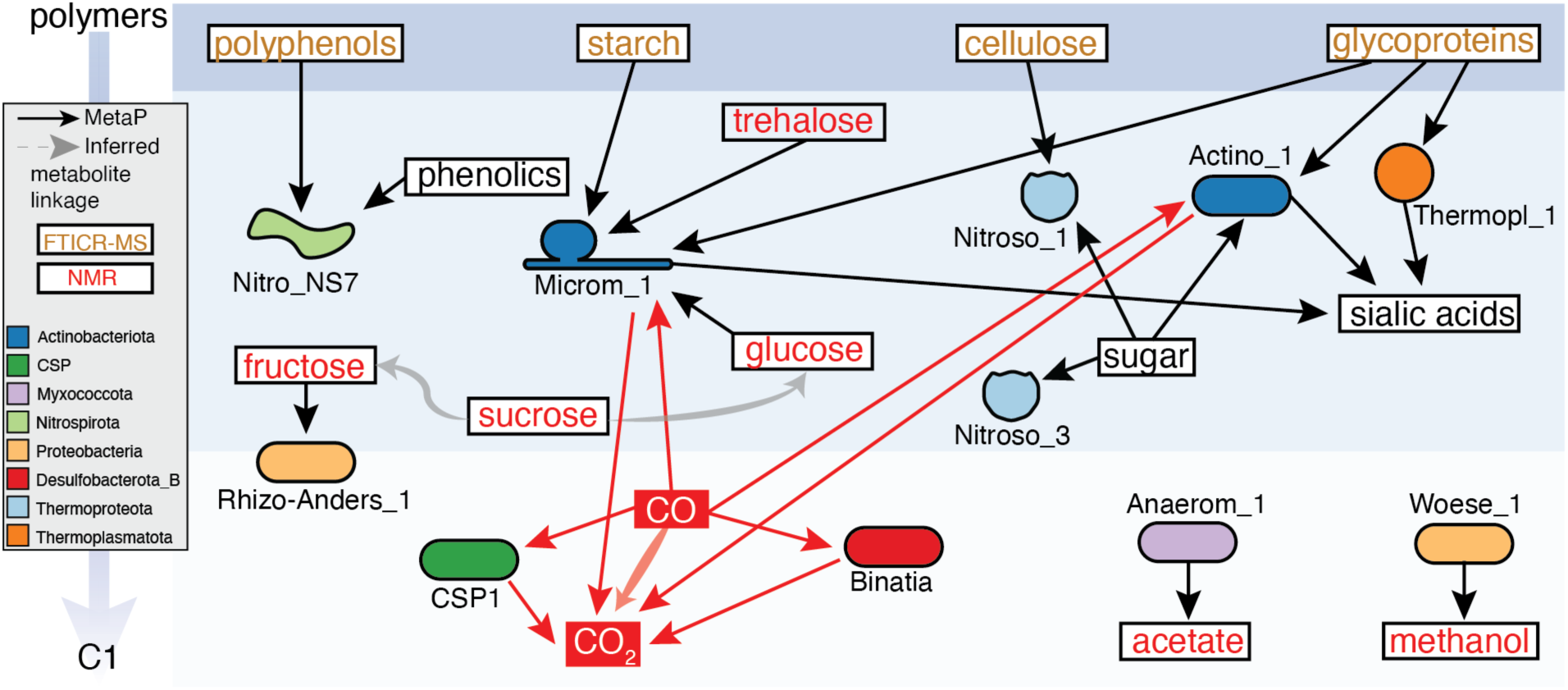
Metaproteomics and metabolomics reveal microbial metabolic handoffs that support carbon cycling in river sediments. Detected metabolites are given in boxes, with NMR-detected compounds listed in red, polymers from FTICR-MS in orange, undetected metabolites in black. These polymers were inferred from FTICR-MS assigned biochemical classes and the specificity of CAZymes detected in metaproteome, where starch and cellulose were within the “polysaccharide-like” class and glycoproteins were in the “amino sugar-like” class. MAG-resolved metaproteome information is indicated by solid arrows, with MAG shape colored by phylum. Red arrows indicate processes leading to CO2 production, while black arrows indicate other microbial carbon transforming genes expressed in the proteome. Shaded bold arrows indicate chemical connections, where(1) grey indicates a metabolite was detected along with putative downstream products (e.g., sucrose conversion to glucose) but metaproteomic lacked evidence for the transformation or(2) red indicates a metabolite not measured but metaproteomic evidence supported transformation (e.g., CO conversion to CO2).

It is well recognized that heterotrophic oxidation of organic carbon derived in HZ sediments largely contributes to river respiration(2). Despite generally low organic concentrations in our sediments (<10 mg/g), FTICR-MS analysis showed that lignin-like compounds were the most abundant biochemical class detected in our samples suggesting that plant litter was likely source of organic carbon (**Fig. S4**). In support of this, 38% of the HUM-V genomes encoded genes for degradation of phenolic/aromatic monomers, while 11% could degrade the larger, more recalcitrant polyphenolic polymers. In fact, our genome analyses revealed that seven unique genomes constituting a new genus within the uncultivated *Binatia* encoded novel pathways for the decomposition of aromatic compounds from plant biomass (phenylpropionic acid, phenylacetic acid, salicylic acid), and xenobiotics (phthalic acid) (**Fig. S5**).

Gene expression of carbohydrate-active enzymes (CAZymes) also supported the degradation of plant polymers like starch and cellulose. We detected the expression of putative extracellular glucoamylase (GH15) and endo-glucanase (GH5) from an *Actinobacteriota* genome (Microm_1) and the *Nitrososphaeraceae* (Nitroso_2), respectively. The integration of our chemical and biological data revealed that heterotrophic metabolism in these sediments could be maintained, likely by inputs of plant biomass. In support of carbon depolymerization, sugars like glucose, and sucrose were detected by nuclear magnetic resonance (NMR).

We next sought to identify microorganisms that could utilize these sugars and found that members expressed transporters for fructose (Rhizo-Anders_1), glucose (Microm_1), and general sugar uptake (Actino_1, Nitroso_2, Nitroso_3). In support of further decomposition, we detected organic acids (acetate, butyrate, lactate, pyruvate, propionate) and alcohols (ethanol, methanol, isopropanol) by NMR. Similarly, proteomics supported interconversions of these smaller carbon molecules, with the *Myxococcota* (Anaerom_1) expressing genes for aerobic acetate respiration and the archaeal *Woeseia* (Woese_1) respiring methanol. In summary, the chemical scaffolding and overlayed gene expression patterns support an active heterotrophic metabolic network in these HZ, likely driven by plant biomass decomposition.

In addition to heterotrophy, our proteomics data revealed autotrophy might also generate carbon dioxide (CO2). Dehydrogenase genes for the aerobic oxidation of carbon monoxide (CO) were among the most prevalent across these sediments. This metabolism was expressed by phylogenetically distinct lineages, including members of uncultivated lineages *Binatia* (Binatia_2) and *CSP1-3* (CSP1_3_1), as well as members of *Actinobacteriota* (Actino_1, Microm_1), *Methylomirabilota* (Roku_AR37_2), and *Proteobacteria* (Burk_1, Thioh_1). The wide range of bacteria and archaea that encoded dehydrogenase genes, combined with gene expression data, suggests carbon monoxide oxidation may be an important metabolism for persistence in HZ sediments.

Given these sediments have relatively low total carbon concentrations (**Fig. S6**), we consider it possible that carbon monoxide may act as a supplemental microbial energy and/or carbon source. Based on genome content, we cautiously infer members of *Actinobacteriota* (Microm_1), *Binatia*, and *CSP1-3* may be capable of carboxydotrophy (i.e., using carbon monoxide as sole energy and carbon source), while the *Actinobacteriota* (Actino_1) is a likely carboxydovore (i.e., oxidize carbon monoxide, while requiring organic carbon). While this metabolism is poorly resolved environmentally, recent efforts have shown it is induced by organic carbon starvation to mediate aerobic respiration, thereby enhancing survival in oligotrophic conditions(26). Here we add river sediments to list of aerated environments (e.g. ocean and soils) where this metabolism may acts as sink or regulate the emission of this indirect GHG(27, 28).

Since proteomics indicated heterotrophy and carbon monoxide oxidation could generate carbon dioxide, we next tracked microorganisms that could fix this compound, sequestering its release. Genome analyses revealed four pathways for carbon fixation were encoded by 75% of HUM-V microbial genomes including (i) Calvin-Benson-Bassham cycle, (ii) reductive TCA cycle, (iii) 3-HydroxyPropionate /4-HydroxyButyrate cycle, and (iv) 3-Hydroxypropionate bi-cycle. The two nitrifying lineages were inferred chemolithoautotrophs, with *Nitrososphaeraceae* encoding HydroxyPropionate/4-HydroxyButyrate (3HP/4HB) and the *Nitrospiraceae* encoding the reductive tricarboxylic acid (TCA) cycle. Other phylogenetically diverse lineages, *Acidobacteriota*, *Binatia, CSP1_3, Proteobacteria*, and *Woeseiaceae* encoded redundant fixation pathways.

Our genome and proteome data revealed the prevalence and activity of single carbon metabolism in these sediments. Carbon monoxide and dioxide are likely the primary substrates, as HUM-V only had minimal evidence for methanol oxidation (*Woeseia*), no methanotrophs, and no methanogens. Together our proteogenomic findings hint at the importance of carbon monoxide and carbon dioxide in sustaining microbial metabolism in these aerated, but low, or fluctuating carbon environments. Further work is needed to understand physiochemical factors controlling carbon monoxide oxidation and carbon dioxide fixation activity, and the balance between production (via heterotrophy and carbon monoxide oxidation) and consumption (fixation) on overall river sediment carbon dioxide emissions.

### Ammonium exchange can support coordinated nitrogen mineralization and nitrification pathways

The ratio of total carbon (C) and total nitrogen (N) (e.g., C/N) is a geochemical proxy used to denote the possible microbial metabolisms that can be supported in a habitat(29, 30). Our HZ sediments had C/N ratios with a mean of 6.5 ± 1.1 (maximum 8.4) (**Fig. S6**). Geochemical theory posits that sediments with low C/N ratios (<15) support organic mineralization that yields sufficient ammonium such that heterotrophic bacteria are not N-limited and nitrifying bacteria are able to compete successfully for ammonium enabling nitrification (30-32). Based on our sediment C/N ratios, we hypothesized organic nitrogen mineralization and nitrification co-occurred in these sediments. Here we profiled the microbial substrates (organic nitrogen metabolites, ammonium) and expressed pathways (mineralization and nitrification) to provide biological validation of this established geochemical theory.

To examine the microbial contributions to organic nitrogen mineralization, we examined metaproteomic data for peptidases, genes that mineralize organic nitrogen into amino acids and free ammonium. Peptidase (n=31) gene expression was three times more abundant and prevalent than glycoside hydrolase genes catalyzing organic carbon transformations, hinting at the relative importance of the former. In support of active microbial N mineralization, FT ICR MS revealed that protein-like and amino sugar like organic nitrogen compounds were correlated to high microbial activity(23), while here we show hydrophobic, polar, and hydrophilic amino acids were abundant (more so than sugars) in the H^1^-NMR characterized metabolites (**Fig. S7**).

The expression of peptidases *in situ*, combined with our genome resolution of their hosts, provided a new opportunity to interrogate the resource sharing and competition underpinning nitrogen mineralization. We first noted which microorganisms were expressing extracellular peptidases (inferred from PSORTb(33)), which would be enzymes that shape external organic nitrogen pools in the sediment. We categorized these expressed peptidases as either releasing free amino acids (end terminus cleaving families, e.g., M28) or releasing peptides (endocleaving families, e.g., S08A, M43B, M36, MO4) (**Fig. 4a**). Members of the *Actinobacteriota*, *Binatia, Methylomirabilota*, and *Thermoproteota* were found to express extracellular peptidases and as such, are likely candidates that contribute to sediment N mineralization.

**Fig. 4.**
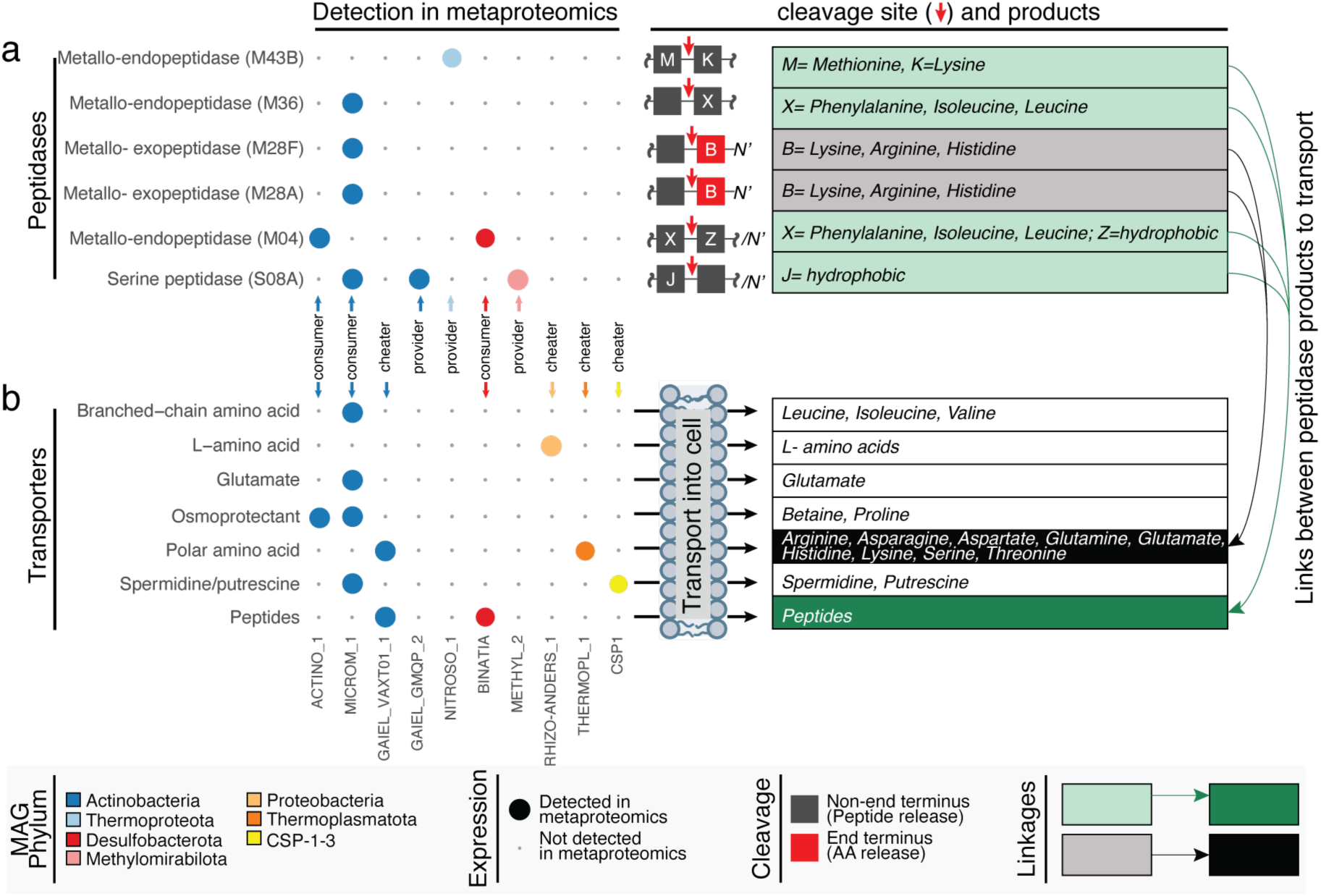
Organic nitrogen mineralization and cellular transport are active microbial processes in river sediments. Bubble plots indicate the expressed genes that were uniquely assigned to specific genome including (**a**) extracellular peptidases and (**b**) cellular transporters for organic nitrogen. Unique peptides detected in at least 3 samples are reported as bubbles and colored by phylum. Table on the right shows putative amino acids cleaved or transported by respective peptidases or transporters, shades of color (green or grey) denote peptides that are cleaved into amino acids that could be transported, providing linkages between extracellular organic nitrogen transformation and transport of nitrogen into the cell. White boxes indicate an organic nitrogen transporter that recruited peptides but could not be linked to outputs of specific peptidases.

We then profiled the expressed amino acid transporters, i.e., genes for the cellular uptake of these smaller organic nitrogen compounds (e.g., branched chain amino acids, glutamate, amines, and peptides were examined) (**Fig. 4b**). Some members functioned exclusively as ‘producers’, expressing peptidases for cleavage of organic N to liberating smaller peptides, yet we could not detect genes for the transport of these produced compounds. Other taxa were ‘producers and consumers’, as genomes in the *Actinobacteriota* and *Binatia* expressed genes for external peptidases and for transporting the organic N products into the cell. Alternatively, we entertain the possibility that members of the *CSP1-3, Proteobacteria*, and *Thermoplasmatota* could be functioning as ‘exploiters’ given that these members expressed genes for assimilating peptidase products but did not contribute to the cost of their production. Expanding on previous carbon degradation frameworks of natural communities(34, 35), we provide a closer examination of the cooperative and competitive interactions that underpin nitrogen mineralization, and further offer new molecular insights into this important terrestrial process.

Finally, we examined the proteomes for evidence that nitrification co-occurred with organic nitrogen mineralization. Supporting this possibility, the substrate ammonium (NH_4_^+^) was detected in all 33 sediment samples (**Fig. S7**). We did not detect genomic evidence for comammox or anammox metabolisms in HUM-V genomes, suggesting aerobic nitrification by metabolic exchanges between organisms drives nitrification in these sediments. Proteomics confirmed aerobic ammonium oxidation to nitrite was performed by Archaeal *Nitrososphaeraceae*. In fact, ammonia monooxygenase (*amo*) subunits were within the top 5% most highly expressed functional proteins in this dataset. The next step in nitrification, nitrite oxidation to nitrate, was inferred from nitrite oxidoreductase (*nxr*) expressed by members of *Nitrospiraceae*. Demonstrating that new lineages first discovered in HUM-V could shape in situ biogeochemistry, we confirmed that 5 genomes from two new species of *Nitrospiraceae* expressed nitrification genes (Nitro_40CM-3_1, Nitro_NS7_3, Nitro_NS7_4, Nitro_NS7_5, and Nitro_NS7_14).

The proteome supported archaeal-bacterial nitrifying mutualism outlined here appear well adapted to the low nutrient conditions present in many HZ sediments, warranting future research on the universal variables that constrain nitrification rates (i.e., ammonium availability, dissolved oxygen, pH) and their role in driving nitrogen fluxes from these systems(36). In conclusion, our microbial data supports that nitrification is concomitant with mineralization in these samples, providing biological evidence to substantiate inferences made from the C/N ratio of these sediments.

### Metabolic handoffs between phylogenetically distinct microorganisms sustain denitrification

Our proteomics suggests that aerobic nitrification could complement allochthonous nitrate from groundwater discharges, contributing to measured nitrate concentrations in excess of 20 mg/L(2, 37). HUM-V genomes with the capacity for nitrate reduction were phylogenetically diverse, with *NarG* or *NapX* encoded in 11 genomes from the *Actinobacteriota*, *Binatia*, *Myxococcota*, and *Proteobacteria,* yet unique peptides were only detected from *Binatia NarG*. Based on gene expression data, nitrite was likely reduced to nitric oxide both by denitrifying *Burkholderia* and nitrifying *Nitrososphaeraceae,* with the metabolic rationale in the archaeal ammonia oxidizers including a detoxification mechanism or generating electron source for ammonia oxidation (38-40).

We did not detect proteomic evidence for nitrous oxide production but did detect *nos* gene expression for reducing nitrous oxide to nitrogen gas. Here the non-denitrifying *Desulfobacterota_D* (Desulf_UBA2774_1, formerly *Dadabacteria*(41)) expressed the *nos* gene for reducing nitrous oxide to nitrogen gas. Phylogenetic analysis revealed this sequence was a “Clade II” *nosZ* sequence, an atypical variant adapted for lower, or atmospheric nitrous oxide concentrations(42). Our activity data adds to emerging interest on these non-denitrifying clade II *nosZ* microorganisms in terrestrial systems, as these microorganisms increase nitrous oxide sink capacity (without contributing to its production)(43, 44). Notably, the activity of this enzyme would have been missed using traditional *nosZ* primers, denoting the value of our untargeted, expression-based approach(45).

We do note that HUM-V capacity for nitrogen cycling exists beyond the proteomics detected instances highlighted above. For example, *Binatia* encoded dissimilatory nitrite reduction to ammonium (DNRA) and the potential for nitrous oxide production via *nor* was encoded by two *Gammaproteobacteria* (Steroid-FEN-1191_1, Steroid_1) and a member of the *Myxococcota* (Anaerom_1). HZ are characterized by dynamic changes in flow direction that lead to cyclic changes in aqueous compositions of dissolved oxygen, nitrate, and organic carbon, thus it is possible that this broader metabolic potential is manifested under other temporal conditions(46).

A key finding of our denitrification metabolic inventory is that complete denitrification by a single microorganism is likely the exception rather than the rule in natural systems(47, 48). In support of this, none of the genomes reconstructed here encoded a complete denitrification pathway for reducing nitrate to nitrous oxide or dinitrogen. Similarly, the metaproteomics data indicated that separate microbial members catalyzed each step of denitrification. Since cross-organism nitrogen chemical exchanges are necessary to support active denitrification, one must consider that physical processes (e.g., advection, diffusion) or the spatial colocalization of microorganisms, as well as organic carbon availability, may have disproportionate impacts on flux of nitrous oxide and dinitrogen in HZ sediments.

### Viral influence on sediment carbon and nitrogen cycling

We reconstructed 2,482 vMAGs that dereplicated into 111 dereplicated viral populations (>10kb) in HUM-V (**Fig. 5a**), making this one of only a handful of genome-resolved studies that include viruses derived from rivers (16, 49, 50). To our knowledge, this is the first study to provide a coordinated analyses of microbial and viral community genomes, and given their sparse sampling, only 5 of the HUM-V viral genomes had taxonomic assignments using viral taxonomies established from standard reference databases. To better understand if the remaining viral genomes had been previously detected in other virally sampled freshwater systems, we compared the protein content of vMAGs in our system to an additional 1,861 viral genomes we reconstructed *de novo* or obtained from public metagenomes from North and South America (**Fig. 5b**). Of the 105 remaining viral genomes, 15% (n=17) clustered with these freshwater derived viral genomic sequences indicating possible cosmopolitan viruses. Of the remaining viral genomes, 23% (n=26) clustered only with genomes recovered in this data set, indicating multiple samplings of the same virus across different sites and depths. The remaining 57% (n=63) of the viral genomes were singletons, meaning they were sampled from these sediments once. Combined, these results hint at the possible biogeographically diverse, as well as endemic viral lineages warranting further exploration in river sediments.

**Fig. 5.**
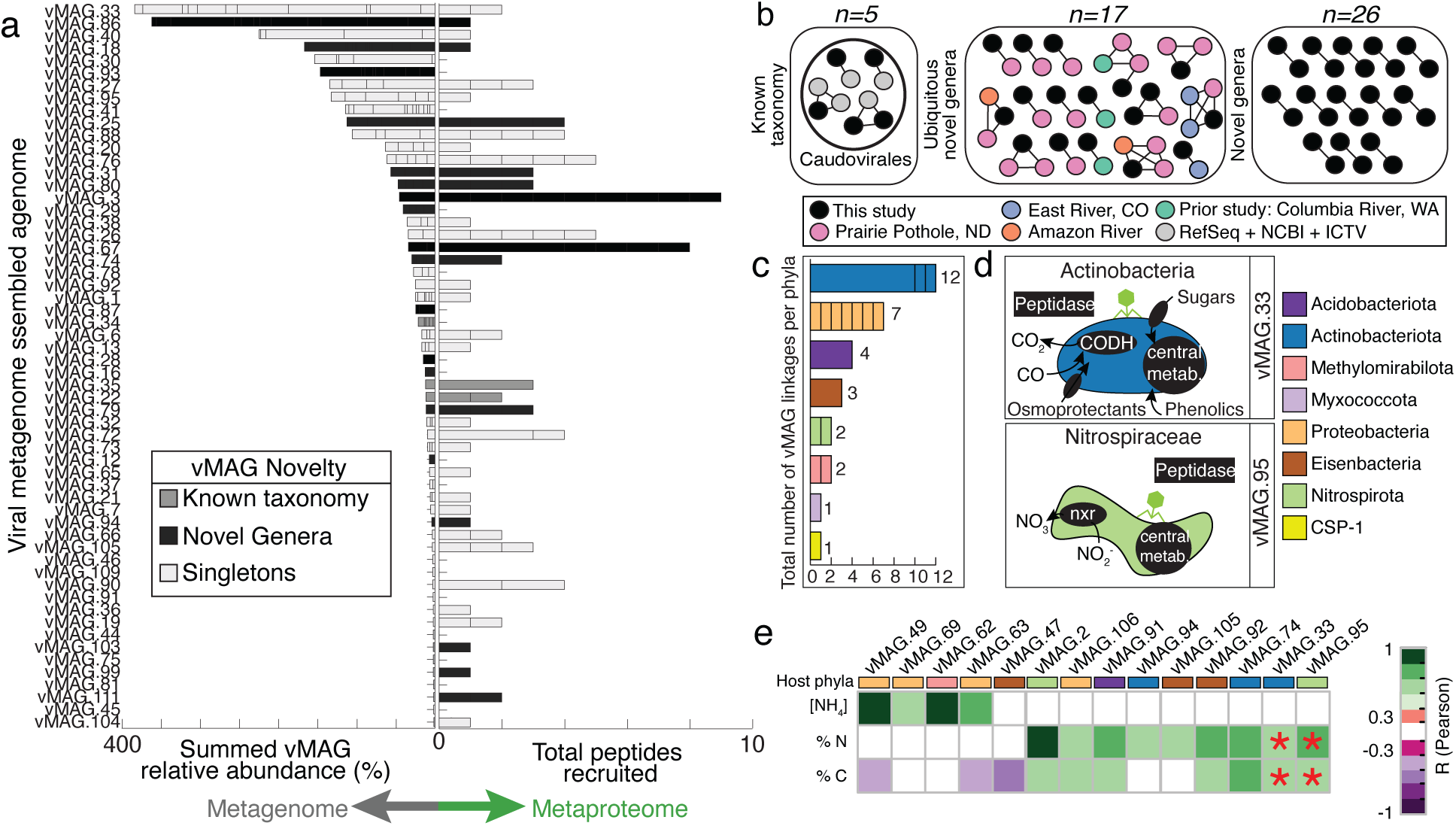
Viruses in HUM-V are active, taxonomically novel, and can play key roles in microbial host metabolism and river geochemistry. **a)** Butterfly plot showing summed genomic relative abundance (left side) and total peptides recruited for each vMAG population (total 111, 58 shown). Bars are colored by clustering of vMAGs from this study with (i) viruses of known taxonomy in RefSeq, ICTV and NCBI Taxonomy (dark grey), (ii) novel genera, both only from this study and ubiquitous (black), and no clustering from any database (light grey, singletons). b) Similarity network of the few vMAGs from our study (black) that clustered to viruses belonging to the default RefSeq, ICTV and NCBI Taxonomy databases (gray), as well as clustering of our vMAGs to other freshwater, publicly available dataset we mined (Pink, Purple, Orange, and Turquoise). The remaining clusters of viruses that were novel (e.g., did not cluster with prior viral genomes) are shown, with the full network file including singletons shown in Table S5. c) Stacked bar chart of the total number of vMAGs (n=32) that have putative host linkages. Each bar represents a phylum and lines within bars indicate the linkages for specific genomes within each phylum. For example, there are three genomes within the Actinobacteriota phylum that collectively have 12 viral linkages and of the three genomes that have linkages, one host has 10 viruses linked, while the other two hosts have 1 virus linked. d) Genome cartoons of microbial metabolisms for two representative genomes that could be predated by vMAGs, with the genes shown in black text boxes denoting processes detected in proteomics. These two microorganisms were selected as examples because they were active members in shaping carbon and nitrogen metabolism in these river sediments but could be impacted by viral predation; other virus-host relationships are reported in Supplementary Figure 9. e) Heatmap reports correlations between a subset of vMAGs with rectangle colors denoting the putative phyla for the respective host. Correlations between these vMAGs and ecosystem geochemistry (NH_4_ µg/gram, %N, %C) are reported with significant correlation coefficients denoted by purple-green shading according to the legend. Red asterisks (*) indicate the vMAG relative abundance predicted a key environmental variable by sparse partial least squares (sPLS) regression. Note two of these predicted vMAGs are shown in (d).

We then assessed peptide recruitment to the viral portion of HUM-V (**Fig. 5a**). For viruses and microbes alike, the most abundant genomes did not have the highest gene expression. While the viral gene expression was not structured by edaphic or spatial factors (**Fig. S8ab**), it was strongly coordinated to the microbial patterns (**Fig. S8c**). Like our microbial genome peptide recruitment, 66% of the viral genomes uniquely recruited peptides. This exceeded prior viral metaproteome recruitment from other environmental systems (e.g., wastewater, saliva, rumen, with ranges from 0.4-15% (35, 51, 52). From this we infer a relatively large portion of the viral community was active at the time of sampling.

The proteomic recruitment of viruses sampled in HUM-V hinted at the possibility that viruses could structure the microbial biogeochemistry through predation. *In silico* analysis assigned a putative host to 29% of the 111 viral genomes. Viruses were linked to 18 microbial genomes that belong to bacterial members in *Acidobacteriota*, *Actinobacteriota*, *CSP1-3*, *Eisenbacteria, Methylomirabilota*, *Myxococcota*, *Nitrospirota*, and *Proteobacteria* (**Fig. S9ab**). Analysis of the metaproteomes for these putative phage-impacted genomes revealed these hosts expressed genes for carbon monoxide oxidation (*Actinobacteriota*), carbon fixation (*CSP1-3*), nitrogen mineralization (*Acidobacteriota, and Methylomirabilota*), methanol respiration (*Myxococcota*), nitrification (*Nitrospirota*), and ammonia oxidation (*Proteobacteria*) (**Fig. 5cd**). Thus, viral predation in HZs could govern carbon and nitrogen biogeochemistry and may explain some of the strain and functional redundancy we observe encoded and expressed by microbes in these sediments.

We next inventoried HUM-V viral genomes for auxiliary metabolic genes (AMGs) with the potential to augment biogeochemistry. We detected 14 auxiliary metabolic genes (AMGs) which we confirmed were not bacterial in origin and had viral-like genes on both flanks(53). These highly ranked AMGs (see Methods) had the potential to augment carbon (CAZymes), sulfur (sulfate adenylyltransferase), and nitrogen (amidase to cleave ammonium) metabolism (**Fig. S9c**). One of our viral genomes that was putatively linked to a *Steroidobacteraceae* (Steroid_1) host genome encoded a pectin lyase gene (PL1). This viral PL1 could enable its host to cleave pectin, generating pectin oligosaccharides that could be used via two host-encoded glycoside hydrolases (GH4 and GH2), ultimately freeing galactose for energy metabolism (**Fig. S9de**). While theoretical, we include this as an example to illustrate how virally encoded genes could expand the substrate ranges for their hosts and alter biogeochemical cycling in river sediments.

In support of their importance to modulating microbial activity and sediment biogeochemistry, we noted that adding the viral genome abundance patterns to microbial genome abundance patterns improved our predictions of river sediment carbon and nitrogen concentrations (**Fig. S10**). In summary, these results indicate that viral predation and AMGs may contribute to river sediment biogeochemistry, either through top down or bottom-up controls on the microbial community. Further interrogation of river sediment viral communities through genomics will yield a more comprehensive understanding of the ecological networks underpinning river biogeochemistry.

## Conclusions

### A multi-omics informed roadmap of carbon and nitrogen cycling in hyporheic zone sediments

Despite the importance of the HZ and its relative accessibility in terms of sampling locations, HZ microbial and viral communities are surprisingly under sampled in a genomic context. Previous studies pertaining to this ecosystem are not genome resolved and used 16S rRNA amplicons or unbinned metagenomes (8–11, 19–21), thus limiting the predictive and explanative power of the study and often times not discriminating between metabolically active and inactive organisms. Further, the few studies which are genome-resolved(21, 22) often focus on specific lineages or single processes and not the entire microbial community, missing the complex interplays between aspects of carbon and nitrogen cycles.

Here we created HUM-V, a genome resolved database to expand on prior non-genome resolved analyses done in a previous publication by our team(23). In concordance to the findings by Graham et al, we failed to detect any significant differences across spatial gradients in community composition or proteomics data, supporting the notion that microbial phenotypic plasticity is important in these sediments. However, our new resolution provided insights into this metabolic plasticity and its mechanistic underpinnings, as we recovered strains with overlapping metabolic potentials from different organisms. Beyond metabolic potentials, our genome-resolved proteomic data further highlighted processes with overlapping expressed genes and provided some of the first activity measurements for members of hyporheic microbiomes. As an example, we focus on new insights gleaned from the 7 recovered *Binatia* genomes (one which included a complete 16S rRNA gene), which are geographically widespread, uncultured, and have only been described in terms of metabolic potential by one previous publication(54). Proteomics demonstrated these bacteria (i) aerobically oxidized carbon monoxide, (ii) mineralized organic nitrogen, and (iii) denitrified, vastly contributing to carbon and nitrogen cycling in these sediments and being biogeographically dispersed (**Fig. S11a**). We also highlighted the metabolic capacity of this lineage in polyphenolic metabolism, perhaps explaining the prevalence of this lineage in terrestrial, plant impacted systems (**Fig. S11b**). These findings illustrate the power of HUM-V in illuminating new roles for members of uncultivated, previously enigmatic microbial lineages in hyporheic zone biogeochemistry.

Empowered by our genome-resolved, process-based metaproteomic analyses (**Fig. 2-Fig. 5**), we present a conceptual model outlining pathways of co-occurring microbial and viral contributions to carbon and nitrogen biogeochemistry in these sediments (**Fig. 6**). Denoted by black arrows, we highlight different microbial transformations that we discovered in our proteomics data. Building on those, colored arrows show how these contributions were complemented by inputs from aquatic sources such as surface or ground water (blue arrows), terrestrial / biotic sources (purple arrows) or atmospheric sources (green arrows). Together, we demonstrated how these could result in the formation and depletion of nutrients in shared resources pools. We further revealed that modes of competition and cooperation form a network of metabolic cross feeding that affected organic and inorganic carbon cycling, and is intertwined with nitrogen mineralization, nitrification, and denitrification pathways.

**Fig. 6.**
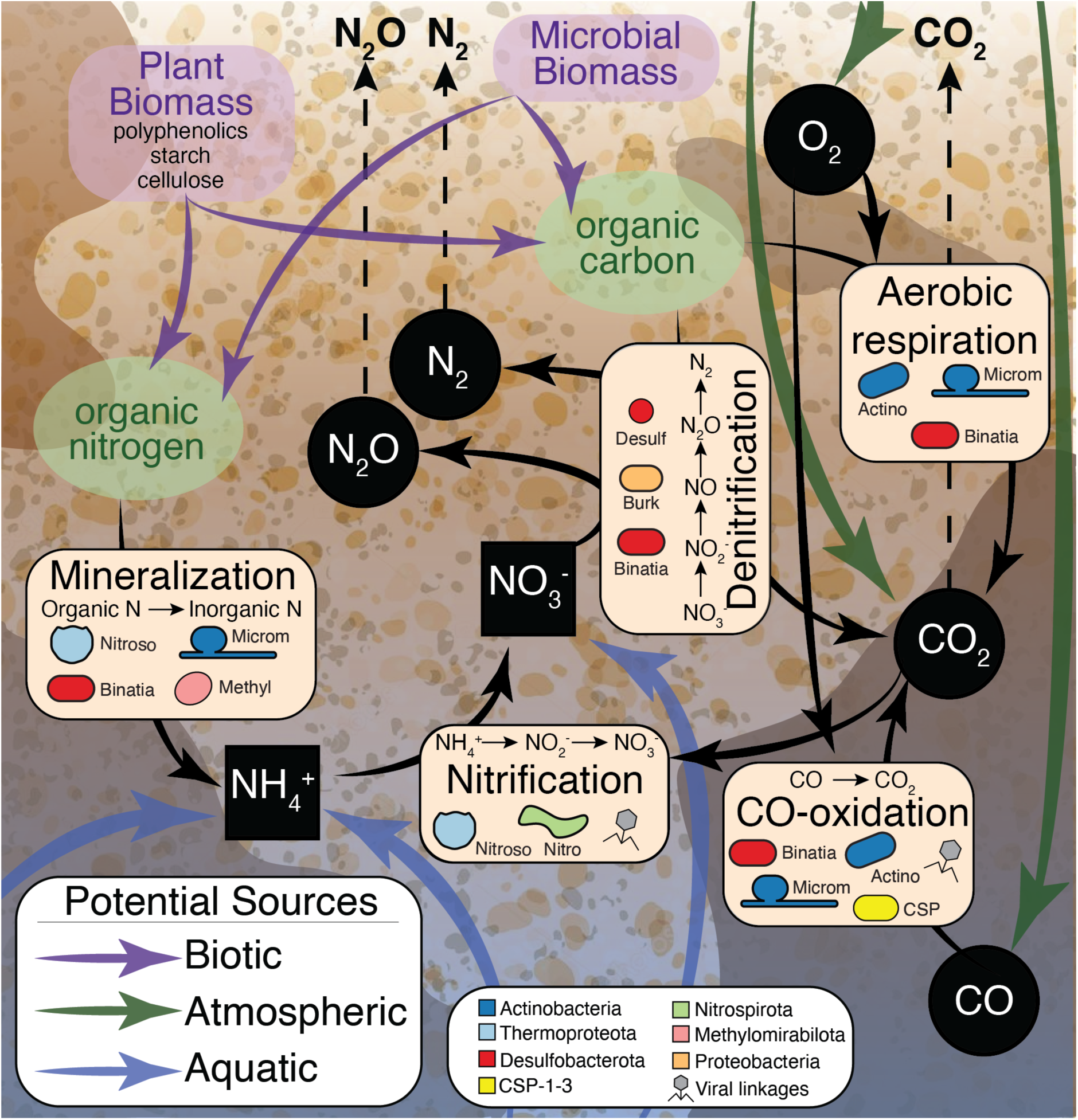
Conceptual model uncovering microbes and processes contributing to carbon and nitrogen cycling in river sediments. Integration of multi-omic data uncovered the microbial and viral effects on carbon and nitrogen cycling in river sediments. Black arrows signify microbial transformations uncovered in our metaproteomic data. Specific processes (e.g., mineralization, nitrification, CO-oxidation, denitrification, and aerobic respiration) are highlighted in beige boxes, with microorganisms inferred to carry out the specific process denoted by overlaid cell shapes colored by phylum. Prior to this research, little was known about the specific enzymes and organisms responsible for river organic nitrogen mineralization and CO-oxidation, thus this research adds new content to microbial roles in carbon and nitrogen transformations in these systems. Possible (biotic, atmospheric, and aquatic) carbon and nitrogen sources are shown by purple, green, and blue arrows respectively. Inorganic carbon and nitrogen sources are shown by black squares (aqueous) and black circles (gaseous) with white text and dashed arrows indicating possible gasses that could be released to the atmosphere. Processes that could be impacted by viruses are marked with grey viral symbols.

In conclusion, while river carbon and nitrogen budgets are often quantified by direct measurements of inputs, and the concentration of inorganic and organic compounds exported from rivers, our findings put forth an integrated framework that advances microbial roles in hyporheic carbon and nitrogen transformations. It yields insights that could inform research strategies to reduce existing predictive uncertainties in river corridor models and resolves microbial contributions and ecological handoffs that were thought to occur but were poorly defined in river sediments. We also highlight previously enigmatic processes that could directly impact river GHG fluxes in unappreciated ways (e.g., carbon dioxide fixation, type II nitrous oxide reduction). Ultimately, we show that a genome-resolved database allows us to track the fates of resource pools of carbon dioxide, ammonium, and inorganic nitrogen to show how consumption and production of these compounds contributes to overall GHG fluxes.

## Materials and Methods

### Sample collection, DNA isolation, and chemical characterization

Samples were collected from the hyporheic zone of the Columbia River (46°22’15.80″N, 119°16’31.52″W) in March 2015 as previously described(23). Briefly, liquid nitrogen frozen sediment profiles (0-60cm) were collected along two transects separated by approximately 170 meters (**Fig. 1abc**). At each transect, three sediment cores up to 60 cm in depth were collected at 5-meter intervals perpendicular to the river flow. Each core was sectioned into 10 cm segments from 0-60-centimeter depths for downstream analyses.

DNA isolation was carried out as previously described(23), with sequencing sent to the Joint Genome Institute (JGI, n=33) under proposal 1781(23). New deep sequencing described here was performed at the Genomics Shared Resource facility at The Ohio State University (OSU, n=10) using a Nextera XT library System. Libraries at both facilities were sequenced using an Illumina HiSeq 2500 platform. **Table S1** details all sequencing information, including accession numbers.

Chemical analyses included geochemical and metabolite data, where geochemistry and Fourier-transform ion cyclotron resonance mass spectrometry (FTICR-MS) methods were performed as previously described(23). Additional metabolite data was obtained through ^1^H Nuclear Magnetic Resonance (NMR) spectroscopy on sediment pore water. Sediment samples were mixed with 200, 300, or 600 μL of MilliQ water depending on the sediment mass (**Table S2**) and centrifuged to remove the sediment. Supernatant (180 µL) was then diluted by 10% (vol/vol) with 5 mM 2,2-dimethyl-2-silapentane-5-sulfonate-*d*6 as an internal standard. All NMR spectra were collected using a Varian Direct Drive 600-MHz NMR spectrometer equipped with a 5-mm triple resonance salt-tolerant cold probe. Chemical shifts were referenced to the 1H or 13C methyl signal in DSS-d6 at 0 ppm. The 1D ^1^H NMR spectra of all samples were processed, assigned, and analyzed using Chenomx NMR Suite 8.3 with quantification based on spectral intensities relative to the internal standard as described previously(55, 56). All geochemical data and methods can be found in **Text S1, Table S2,** and **Table S3**.

### Metagenome assembly and binning

Raw reads were trimmed for length and quality using Sickle v1.33 (https://github.com/najoshi/sickle) and then subsequently assembled using IDBA-UD 1.1.0(57) with an initial kmer of 40 or metagenomic metaSPAdes 3.13.0(58) with default parameters. To further increase genomic recovery, for the ten samples that had shallow and deep sequencing, metagenomic reads were coassembled using IDBA-UD 1.1.0 with an initial kmer of 40. All assemblies, including co-assemblies, were then individually binned using Metabat2(59) with default parameters to obtain MAGs.

For each bin, genome completion was estimated based on the presence of core gene sets using Amphora2(60). Bins were discarded for further analysis if completion was <70% or contamination was >10% to select for only medium to high quality bins(24). This resulted in 102 MAGs that were then dereplicated using dRep(61) with default parameters and resulted in a final set of 55 MAGs (>99% ANI). To further assess bin quality, we used the Distilled and Refined Annotation of MAGs (DRAM)(53) to identify ribosomal ribonucleic acids (rRNAs) and transfer ribonucleic acids (tRNAs). Genome quality information reported in **Table S4**.

### Metabolic analysis of MAGs

Medium and high-quality MAGs were taxonomically classified using the Genome Taxonomy Database (GTDB) Toolkit v1.3.0 on September 2020(62). Novel taxonomy was identified as the first taxonomic level with no designation using GTDB taxonomy. MAG scaffolds were annotated using the DRAM pipeline(53). Phylogenetic analysis was performed on genes annotated as respiratory nitrate reductase (*nar*) and nitrite oxidoreductase (*nxr*) to resolve novel *Binatia* role in nitrogen cycling. Specifically, sequences from(63) were downloaded and combined with *nar* and *nxr* amino acid sequences from dereplicated bins, aligned using MUSCLE, version 3.8.31, and run through ProtPipeliner, a Python script developed in-house for generation of phylogenetic trees (https://github.com/TheWrightonLab). Phylogenetic trees are provided in Zenodo here: https://doi.org/10.5281/zenodo.6339808. For polyphenol and organic polymer degradation, we used functional annotation in addition to predicted secretion to assess functional potential. To determine if the predicted genes encoded a secreted protein, we used pSortb(33) and SignalP(64) to predict location; if those methods did not detect a signal peptide, the amino acid sequence was queried to SecretomeP and a SecP score > 0.5(65) was used as a threshold to report non-canonical secretion signals. Metabolic information for each MAG discussed in this manuscript are available in **Table S4.**

### Viral Analyses

Metagenomic assemblies (n=43) were screened for DNA viral sequences using VirSorter v1.0.3 with the ViromeDB database option(66), retaining viral contigs ranked 1, 2, 4 or 5 where category 1-2 indicate high confidence predicted lytic viruses and 4-5 indicate high-confidence prophage sequences from VirSorter output(66). Viral sequences were filtered based on size to retain those greater than or equal to 10kb based on current standards(67). Viral scaffolds were then clustered into vMAGs at 95% ANI across 85% of the shortest contig using ClusterGenomes 5.1 (https://github.com/simroux/ClusterGenome)(67). After clustering, vMAGs were manually confirmed to be viral by looking at DRAM-v annotations and further assessing the total viral-like genes relative to non-viral genes, where vMAGs containing more than 18% of non-viral genes were deemed suspicious, manually confirmed non-viral, and subsequently discarded (J flag, DRAM-v)(53).

To determine taxonomic affiliation, vMAGs were clustered to viruses belonging to viral reference taxonomy databases NCBI Bacterial and Archaeal Viral RefSeq V85 with the International Committee on Taxonomy of Viruses (ICTV) and NCBI Taxonomy using the network-based protein classification software vContact2 v0.9.8 using default methods(68, 69). To determine geographic distribution of viruses in freshwater ecosystems, we included viruses mined from publicly available freshwater metagenomes in vContact2 analyses: 1) East River, CO (PRJNA579838) 2) A previous study from the Columbia River, WA (PRJNA375338) 3) Prairie Potholes, ND (PRJNA365086) and 4) the Amazon River (PRJNA237344). The viral sequences that were identified from these systems and the genes used for vContact2 are deposited on Zenodo with doi 10.5281/zenodo.6310084.

Viral contigs were annotated with DRAM-v(53). Genes that were identified by DRAM-v as being high-confidence possible auxiliary metabolic genes (auxiliary scores 1-3)(53) were subjected to protein modeling using Protein Homology / AnalogY Recognition Engine (PHYRE2)(70). Auxiliary scores were assigned by DRAM(53), based on the following ranking system: A gene is given an auxiliary score of 1 if there is at least one hallmark gene on both the left and right flanks, indicating the gene is likely viral. An auxiliary score of 2 is assigned when the gene has a viral hallmark gene on one flank and a viral-like gene on the other flank. An auxiliary score of 3 is assigned to genes that have a viral-like gene on both flanks. To identify likely vMAG hosts, oligonucleotide frequencies between virus (n=111) and non-dereplicated hosts (n=102) were analyzed using VirHostMatcher using a threshold of d2* measurements of <0.25(71). The lowest d2* value for each viral contig <0.25 was used. All vMAG data is reported in **Table S5**.

### Genome relative abundance calculations and their use in predictions

To estimate the relative abundance of each MAG and vMAG, all metagenomic reads for each sample were rarified to 3Gbp and mapped to 55 unique MAGs via Bowtie2 (72) (55, 73). For MAGs, a minimum scaffold coverage of 75% and depth of 3x required for read recruitment at 7 mismatches. For vMAGs, reads were mapped using Bowtie2(72) at a maximum mismatch of 15, a minimum contig coverage of 75% and a minimum depth coverage of 2x. Relative abundances for each MAG and vMAG were calculated as their coverage proportion from the sum of the whole coverage of all bins for each set of metagenomic reads. Genome relative abundances per sample for MAGs and vMAGs are reported in **Table S4** and **Table S5**. Correlations and sparse Partial Least Squares Regression (sPLS) predictions (PLS R package(74)) were done using mapping data pertaining to only the 10 deeply sequenced metagenomes rarified to 4.8Gbp (**Table S6**).

### Metaproteome generation and peptide mapping

Sediment samples were prepared for metaproteome analysis as previously reported in Graham et al. 2018(23) and the protocol outlined by Nicora et al(75). As previously described(55, 76), tandem mass spectrometry (MS/MS) spectra from all liquid chromatography tandem mass spectrometry (LC-MS/MS) datasets were converted to ASCII text (.dta format) using MSConvert (http://proteowizard.sourceforge.net/tools/msconvert.html) and the data files were then interrogated via target-decoy approach(77) using MSGF+(78). For protein identification, spectra were searched against two files that included (i) 55 dereplicated MAG and (ii) 111 clustered vMAGs amino acid sequences. See **Text S1** for more details on metaproteome analysis. Metaproteomic mapping results for MAGs and vMAGs can be found on **Table S7**.

## Data availability

The datasets supporting the conclusions of this article are publicly available. Sequencing data are available in NCBI under BioProject PRJNA576070, with MAGs deposited under Biosamples SAMN18867633-SAMN18867734 and 16S rRNA amplicon sequences under accession numbers SRX9312157-SRX9312180. The raw annotations for each genome are deposited on Zenodo with the following DOI: https://doi.org/10.5281/zenodo.5128772. 111 vMAGs have been deposited NCBI under the BioProject ID PRJNA576070 and in Zenodo within the following DOI: https://doi.org/10.5281/zenodo.5124937. Additionally, the dataset of freshwater viruses we used to cluster to the HUM-V viruses is also hosted on Zenodo with DOI https://doi.org/10.5281/zenodo.6310084. The raw annotations for each viral genome are on **Table S5**. Metaproteomics data are deposited in the MassIVE database under accession MSV000087330. Metabolomics data are publicly available and deposited in Zenodo doi https://doi.org/10.5281/zenodo.5076253. All scripts used in this manuscript are available at https://github.com/WrightonLabCSU/columbia_river.

## Acknowledgements

This work was supported by the Subsurface Biogeochemical Research (SBR) program (DE-SC0018170); the National Sciences Foundation Division of Biological Infrastructure [#1759874]; and the U.S. Department of Energy, Office of Science, Office of Biological and Environmental Research, Environmental System Science (ESS) program through subcontract from the River Corridor Scientific Focus Area project at Pacific Northwest National Laboratory. J.R.R. was partially supported by the National Science Foundation (NRT-DESE) [1450032], A Trans-Disciplinary Graduate Training Program in Biosensing and Computational Biology at Colorado State University. The NMR data, FTICR-MS data and MS-proteomics data in this work was collected using instrumentation in the Environmental Molecular Science Laboratory (grid.436923.9), a DOE Office of Science User Facility sponsored by the Office of Biological and Environmental Research and located at Pacific Northwest National Laboratory. Pacific Northwest National Lab is operated by Battelle for the DOE under Contract DE-AC05-76RL01830. Metagenomic sequencing for this research was performed by the Joint Genome Institute via a large-scale sequencing award (Award 1781) and at the Genomics Shared Resource Core at The Ohio State University Comprehensive Cancer Center supported by P30 CA016058.

The authors would like to thank Tyson Claffey and Richard Wolfe for Colorado State University server management; Sandy Shew for management of computing resources retained from The Ohio State University Unity cluster; Dr. Pearlly Yan at the Genomics Shared Resource Core at The Ohio State University Comprehensive Cancer Center for management of metagenomic sequencing; and Dr. J John for the continuous support. The authors declare they have no competing interests. J.R.R. and M.A.B. contributed equally to this work.

## Supplementary Figure Legends

**Figure S1: DRAM annotation of MAGs.** Heatmap showing the DRAM product output for medium and high-quality genomes (n=102) from HUM-V. The interactive version of this heatmap is available here: https://zenodo.org/record/5124964

**Figure S2: Resazurin reduction assay measurements.** Box and whisker plots showing the values reported for a resazurin reduction assay. Different boxes represent the different depths (0-60cm) that were sampled per sediment core.

**Figure S3: NMDS and Venn Diagram of differences in transect (N / S) or Depth (0-60cm) for MAG proteome. a)** NMDS of north (green) and south (orange) transects of recruited proteome peptides for MAGs. ANOSIM is reported. **b)** NMDS of sediment core depth (0-60cm) of recruited proteome peptides for MAGs. ANOSIM is reported. **b)** Euler diagram showing the number of total proteins recruiting peptides in each transect, where the overlap represents the proportion of these proteins recruiting peptides in both transects. Only proteins recruiting 2 or more total peptides were included (n=898) to allow for recruitment of at least 1 peptide for a given protein to both transects or both depths and reduce false positives.

**Figure S4: Relative abundance of FTICR-MS classes across samples.** The relative abundance of biochemical classes identified in FTICR-MS across samples is not statistically structured by depth. Bar plots show the average relative abundance for each class, with error bars representing one standard deviation (n=33). Individual data points are plotted for each sample with point size increasing with depth. Peaks were classified as described in **Methods**. Raw data is provided in **Additional File 4.**

**Figure S5: Binatia MAGs possess the ability to metabolize compounds related to carbon and nitrogen cycling.** The seven HUM-V genomes reconstructed here encode diverse pathways for transforming phenolic compounds. Pathways are shown with boxes corresponding to each enzyme in the pathway, colored by the number of Binatia MAGs that encode each step. Gene information shown here is reported in detail in **Additional File 6**.

**Figure S6: C:N ratio and total carbon and nitrogen measurements across depths reveal no significant differences.** Box and whisker plots report concentrations. Graphs show C:N ratio, percent total carbon, and percent total nitrogen measurements across different depths (0-60cm). No significant differences were observed.

**Figure S7: NMR detected metabolites and ammonium indicate prevalence of nitrogen-containing compounds, saccharides, organic acids, and alcohols.** Bar graphs showing specific NMR metabolites and ammonium with the percent of total samples that they were found in. Shading denotes the different categories of detected compounds. For detailed information on how these were collected see methods section. Only metabolites detected in more than 4 samples are shown.

**Figure S8: vMAG expressed peptides are not structured by transect of depth, and relative abundance NMDS of vMAGs is coordinate with relative abundance NMDS of MAGs: a)** NMDS of north (green) and south (orange) transects of recruited proteome peptides for vMAGs. ANOSIM is reported. b) NMDS of sediment core depth (0-60cm) of recruited proteome peptides for vMAGs. ANOSIM is reported. c) Procrustes ordination of MAG and vMAG NMDS ordinations using relative abundance genome data across 33 shallower sequenced samples.

**Figure S9: Virus-Host associations show viruses could infect key players in river sediment carbon and nitrogen cycling. a)** Genome cartoons colored by their phyla. Gray dotted circles represent vMAGs that putatively infect each genome. **b)** A breakdown of each vMAG name for each of the infected host genomes. Colors match genome host phyla assignment. **c)** Bar chart showing the different types of AMGs identified in viral genomes with putative hosts, with the predicted substrate for each AMG denoted by colored boxes on the y-axis. **d)** Viruses encode AMGs which can expand the metabolic repertoire of their microbial hosts. Conceptual models of a putative glycoside hydrolase (GH) and a pectin lyase (PL) are shown. **e)** For one putative host, Steroidobacteraceae, with a linkage to a viral genome that encodes a PL1, a conceptual model of an integrated metabolism is shown. Specifically, integration of vMAG.63 genome into the host genome may provide the capacity to cleave the pectin backbone (PL1), followed by oligo cleavage of pectin, resulting in galactose monomers for host cell metabolism.

**Figure S10: sPLS regressions show vMAG abundance is better predictor of carbon and nitrogen relative to MAGs.** sPLS plots showing predicted vs measured values for percent carbon (c_per) and percent nitrogen (n_per) for our samples using n=10 deep sequences. Shown sPLS are **a)** vMAGs only, **b)** MAGs only, and **c)** vMAGs and MAGs combined. Values corresponding to cor.test function output in R for the predicted and measured values are shown in each box (t, degrees of freedom, p-value, confidence interval, and correlation).

**Figure S11: Uncultured Binatia are widely dispersed across ecosystems and express nitrogen and carbon genes *in situ*. a)** Using the 16S rRNA gene (from Binatia_7), we inventoried the distribution of closely related species to our HUM-V genomes (>97% similarity) in the Sequence Read Archive (SRA) samples, uncovering the ecological distribution of these organisms from soils, as well as a wide variety of terrestrial, terrestrial-aquatic, marine samples, indicating the processes uncovered by proteomics here are likely applicable to a wide range of ecosystems. **b)** Metabolic genome cartoon for the major functions encoded by the Binatia MAGs. Dotted arrows are functions encoded in the metagenome, with black arrows corresponding to metaproteome-detected enzymes. (BCAA, Branched chain amino acids).

## Supplementary Files Index

*Table S1:* Metagenome read counts, accession numbers, biosamples.

*Table S2:* Geochemistry data and NMR metabolite data. *Text S1:* Supplementary materials and methods

*Table S3:* FTICR-MS Data.

*Table S4:* MAGs. Completion information, accession numbers, annotations, Read mapping information, metabolism summary from DRAM.

*Table S5:* vMAGs. MIUViG information, accession numbers, annotations, read mapping information, vContact2 results, downloaded MetaG information for vContact2 analyses, virhostmatcher2 results.

*Table S6:* SPLS data and correlations results

*Table S7:* Metaproteomes. MAG Unique peptides, MAG unique peptide NSAF relative abundance, MAG Per genome NSAF Relative abundance, vMAG unique peptides

